# Scalable image processing techniques for quantitative analysis of volumetric biological images from light-sheet microscopy

**DOI:** 10.1101/576595

**Authors:** Justin Swaney, Lee Kamentsky, Nicholas B Evans, Katherine Xie, Young-Gyun Park, Gabrielle Drummond, Dae Hee Yun, Kwanghun Chung

**Author notes:** Correspondence should be addressed to K.C. These authors contributed equally to this work (Justin Swaney, Lee Kamentsky).

## Abstract

Here we describe an image processing pipeline for quantitative analysis of terabyte-scale volumetric images of SHIELD-processed mouse brains imaged with light-sheet microscopy. The pipeline utilizes open-source packages for destriping, stitching, and atlas alignment that are optimized for parallel processing. The destriping step removes stripe artifacts, corrects uneven illumination, and offers over 100x speed improvements compared to previously reported algorithms. The stitching module builds upon Terastitcher to create a single volumetric image quickly from individual image stacks with parallel processing enabled by default. The atlas alignment module provides an interactive web-based interface that automatically calculates an initial alignment to a reference image which can be manually refined. The atlas alignment module also provides summary statistics of fluorescence for each brain region as well as region segmentations for visualization. The expected runtime of our pipeline on a whole mouse brain hemisphere is 1-2 d depending on the available computational resources and the dataset size.

## Introduction

Light-sheet fluorescence microscopy (LSFM) is an optical sectioning technique that provides high-speed acquisition of high resolution images. Affordable open-access systems have promoted adoption of LSFM^1^. As a result, LSFM has become commonplace in the study of complex biological systems^2–5^. However, the high-throughput acquisition offered by LSFM can quickly generate terabytes of image data, posing challenges in data storage, processing, and visualization. These challenges must be addressed in order to perform the quantitative analyses needed to answer the complex biological questions at hand.

SHIELD is a tissue transformation technique that preserves endogenous biomolecules for imaging within intact biological systems^6^. SHIELD retains fluorescent protein signals through the clearing process and is compatible with stochastic electrotransport staining, allowing visualization and quantification of fluorescence signals throughout the entire brain^7^. When SHIELD-processed tissues are imaged using LSFM, entire organs such as the mouse brain can be imaged at single-cell resolution in just 2 hours, offering more data than was previously available to answer new biological questions.

Here we present detailed protocols for quantifying fluorescence signals in each brain region of SHIELD-processed mouse brain LSFM datasets. The pipeline is composed of modules for image destriping, stitching, and atlas alignment. Each module can either be used independently or in combination to perform region-based statistical analyses of fluorescence within intact mouse brain samples. The pipeline and all its dependencies have been packaged into a single Docker container, allowing for simple installation and cross-platform use. The ease of deployment offered by Docker makes our image processing pipeline more accessible to researchers without much programming experience. We also provide a dataset of a whole mouse brain hemisphere for users to test our pipeline.

### Development of the protocol

In order to analyze large-scale volumetric images acquired using LSFM, research labs typically create their own image processing pipelines^4^,^8–10^. These image processing pipelines are designed to solve specific problems in applying LSFM to the study of complex biological systems. Real-time cell tracking systems have been reported to study the dynamics of embryogenesis in *D. melanogaster*^8,9^. The cell tracking pipeline relies on optimized CUDA programming to achieve real-time performance. Several computational pipelines geared toward processing time-lapse images of *D. melanogaster* assume that each time point image is smaller than the amount of available memory. In contrast, LSFM of whole mammalian organs often generates individual volumetric images that are larger than the amount of available memory.

Recently, LSFM images of a whole mouse brain have been used to create a single-cell mouse brain atlas^4^. The pipeline consisted of a heterogeneous mix of MATLAB, Python, and C++ software as well as expensive computer hardware, including a dedicated image processing server equipped with four NVIDIA graphics processing units (GPU). In order to handle individual volumetric images that are larger than the amount of available memory, images were processed slice-by-slice for cell detection and rescaled to a manageable size for atlas alignment. Although computationally impressive, such tools often require a great deal of programming expertise or access to proprietary software. As a result, the current large-scale image processing pipelines may be inaccessible to non-experts, and there is a need for large-scale image processing tools for researchers focused on biological questions rather than computational challenges.

The protocols presented here are designed to be easy to setup and applicable to users without much experience in setting up complex development environments. Since some users may only want to use part of our pipeline, the protocols are partitioned into three computational modules: first, our image destriping for removing streaks and performing flat-field correction in raw LSFM images; second, stitching for creating a single 3D image from the individual 2D images; and third, semi-automatic atlas alignment for segmenting brain regions and quantifying fluorescence (Fig. 1). Our protocols have been tested on images of SHIELD-processed mouse brain hemispheres acquired with an axially-swept light-sheet microscope.

**Figure 1.**
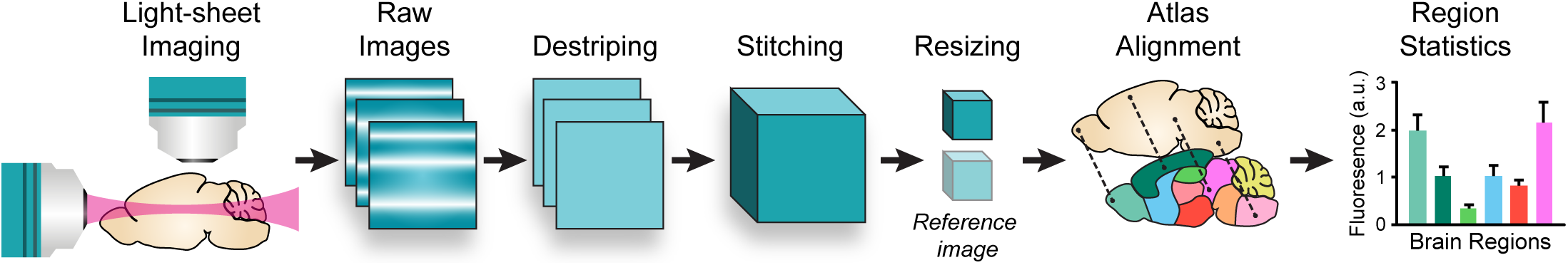
Overall image processing pipeline for whole brain analysis. Raw images from LSFM of a SHIELD-processed mouse hemisphere are destriped and corrected for uneven illumination. Destriped images are then stitched into a multichannel volumetric image, which is resampled to match a reference atlas. Point correspondences from an automatic alignment procedure are manually refined to obtain a region segmentation for the full-resolution, stitched image. The region segmentation is then used to quantify mean fluorescence in each brain region.

#### Development of the destriping module

Stripe artifacts are commonplace in images acquired with LSFM due to irregularities in the refractive index (RI) of the sample^3,11^. This RI mismatch can be compensated for using an immersion medium that has a similar RI to that of the sample^12^. However, the material properties of biological tissues, including the RI, are generally not uniform throughout, making some degree of RI mismatch inevitable. RI mismatch usually results in uneven illumination patterns due to optical aberrations that disrupt the incident light.

Current strategies for image destriping are either based on optical filtering or digital filtering^11,13–15^. Optical filtering strategies attempt to compensate for RI mismatch during imaging, effectively removing the stripe artifacts from the source. However, these methods may disrupt the axial resolution of the LSFM system in the process and may not be applicable to large biological samples. In contrast, digital filtering strategies attempt to remove the stripe artifacts after acquisition by exploiting the noise characteristics induced by the optical aberrations. Since digital destriping methods are implemented as image filters, they can be applied more generally to any images with stripe artifacts.

Previous digital destriping methods have included hybrid wavelet-FFT filters, variational removal of stationary noise (VSNR), and multidirectional filters using the contourlet transform (MDSR)^13–15^. Although VSNR and MDSR have shown superior destriping performance, they are prohibitively slow for applying to whole-brain datasets. The hybrid wavelet-FFT filter is the fastest destriping method of these, but its implementation requires a MATLAB license to use and is single-threaded.

In order to provide a fast, open-source destriping solution, we implemented a new digital destriping tool called pystripe. Pystripe is a Python implementation of the previously reported hybrid wavelet-FFT destriping method with parallel processing support and other improvements. Pystripe uses open-source tools instead of proprietary software such as MATLAB. As in the hybrid wavelet-FFT approach, the amount of filtering in pystripe can be tuned using a bandwidth parameter. Pystripe also adds support for a dual-band filtering mode where the background and foreground of the images can be filtered with separate bandwidths.

#### Development of the stitching module

Imaging large samples with LSFM involves acquiring partially overlapping image stacks which can be stitched together into a single image stack. Several open-source stitching packages are available^16,17^. Terastitcher has been widely adopted for stitching large volumetric images acquired with LSFM. However, the Terastitcher merging step executes within a single thread by default, resulting in longer execution times than necessary. It should be noted that the Terastitcher team provides a parallelized version of Terastitcher based on message passing interface (MPI) upon request, but we found implementing our own merging step based on the multiprocessing module in Python to be more straightforward than managing MPI.

To address these shortcomings, we created the TSV (Terastitcher Volume) module, which implements the Terastitcher merging step in Python with support for lossless TIFF compression and parallel processing. TSV uses the stack displacements computed from Terastitcher to create a memory-mapped array representing the entire image volume. Multiple workers use this memory-mapped array to convert individual images into a single stack, providing faster overall execution. Each worker merges images together and then saves the result using lossless TIFF compression, resulting in lower overall dataset sizes.

#### Development of the atlas alignment module

In order to segment whole-brain LSFM images into different brain regions, the stitched dataset must be registered to a reference atlas, such as the Allen Mouse Brain Atlas (ABA)^18^. The ABA consists of an averaged anatomical reference image of autofluorescence and the corresponding region segmentation image. The ABA also contains tools for registering 3D reconstructions from histological sections to the atlas. However, research labs have resorted to custom atlas alignment methods for LFSM images of intact brain samples^4^.

Elastix is an open-source medical image registration library that is widely used for non-rigid atlas alignment^19^. Elastix performs non-rigid atlas alignment by maximizing the mutual information between source and reference images. Elastix was found to have the highest mutual-information benchmark scores in image registration of cleared brain samples among five freely-available software packages^20^. The global optimization of mutual information is difficult to scale to whole-brain LSFM datasets since the entire dataset cannot be stored in memory. Following previous work on atlas alignment, we address this issue by rescaling the source image to be a similar size compared to the reference atlas, which is a more manageable size. We use the alignment computed from Elastix to generate a set of approximate point correspondences which can manually refined.

To visualize the atlas alignment and edit the approximate point correspondences, we created an interactive web-based registration tool called nuggt (NeUroGlancer Ground Truth). Nuggt is built on an open-source visualization package called Neuroglancer. Nuggt does not modify the underlying Neuroglancer code but rather wraps it into a convenient package for interactive LSFM visualization and atlas alignment. Nuggt displays the source and reference images side-by-side along with the point correspondences overlaid on each image. Using nuggt, the point correspondences can be edited and adjusted to improve the atlas alignment. The source image can also be warped while editing the point correspondences, providing rapid visual feedback of the atlas alignment accuracy.

#### Development of the Docker image

One of the main challenges in adopting a new computational pipeline is obtaining the dependencies and recreating the runtime environment that was intended by the developers. In order to simplify the installation of our pipeline, we packaged our pipeline and all of its dependencies into a single Docker container by creating a Dockerfile describing our runtime environment. Docker provides a consistent, light-weight virtual Linux environment on all major operating systems, and our container has been successfully tested on Windows, Mac, and Linux. Docker can be installed from the Docker website, and our container can be downloaded from Docker Hub. By containerizing our pipeline and using web-based visualization, our modules can either be run locally or on dedicated image processing servers which can be accessed from other clients.

### Applications of the method

The overall image processing pipeline described in this protocol is designed to calculate fluorescence summary statistics from whole-mouse brain images acquired with LSFM on a per-region basis. Our pipeline has been used to quantify mRuby2 and EGFP fluorescence of virally labeled neurons and presynaptic terminals in SHIELD-processed mouse brain hemispheres^6^. Thus, the overall pipeline may be applied in systems neuroscience to quantify fluorescent reporters in cleared samples from mouse models. However, the individual modules that comprise the overall pipeline can also be used independently.

Pystripe can be applied to any images corrupted with horizontal or vertical stripe artifacts. We restricted pystripe to filtering horizontal or vertical stripes because the illumination beam path in most LSFM systems is aligned with the camera detector. Pystripe can, therefore, also be used with multi-view LSFM systems that rotate the sample rather than change the orientation of the illumination beam path^21^. Pystripe also includes the ability to provide a reference flat-field image for illumination correction of vignetting and other stationary artifacts.

TSV can be used to merge an array of partially overlapping image stacks saved in Terastitcher hierarchal format into a single image stack^17^. The memory-mapped array used for stitching is also useful for retrieving sub-volumes of image data. TSV also includes optional utilities for partitioning the stitched image into smaller, uniformly shaped chunks for custom parallel processing. The stitched images can be stored in Neuroglancer precomputed format and served via HTTP, allowing for efficient visualization of whole-brain LSFM datasets at full resolution either locally or over the web.

Nuggt has been used to register SHIELD-processed mouse brain hemisphere datasets to the ABA. Our alignment protocol can be used to register a pair of 3D volumes with mutual information between them and a gradient of mutual information in the initial overlap that can be followed to modify the alignment. In our experience, images that have 90% overlap and rotations of a few degrees can be aligned using this method. The automatic alignment can be refined by adding manually-placed correspondences. The atlas alignment module also provides utilities for calculating image statistics for each brain region given the aligned atlas segmentation.

### Comparison with other methods

Many standard solutions exist for similar tasks addressed in our image processing pipeline. In this section, we compare the methods used in our protocol to existing methods in the context of whole brain LSFM image analysis.

#### Destriping

The previously reported digital destriping algorithm MDSR has achieved state of the art destriping performance on LSFM images^15^. MDSR relies on the contourlet transform to perform energy compaction of striping artifacts in arbitrary orientations, whereas pystripe uses the discrete wavelet transform to remove either horizontal or vertical striping artifacts.

When comparing the resulting images from MDSR and pystripe, similar filtering performances are observed from both methods on our test images (Fig. 2a). This suggests that the contourlet transform does not drastically improve energy compaction of the stripe artifacts compared to the discrete wavelet transform when the stripes are oriented horizontally. When comparing the average execution speed for destriping using MDSR and pystripe on a single core, MDSR takes over 30 min per frame, while pystripe takes only 5 seconds (Fig. 2b). Using multiple cores, the destriping frame rate for pystripe increases linearly with the number of cores, reaching 8 frames per second with 48 cores (Fig. 2c, Supplementary Video 1).

**Figure 2.**
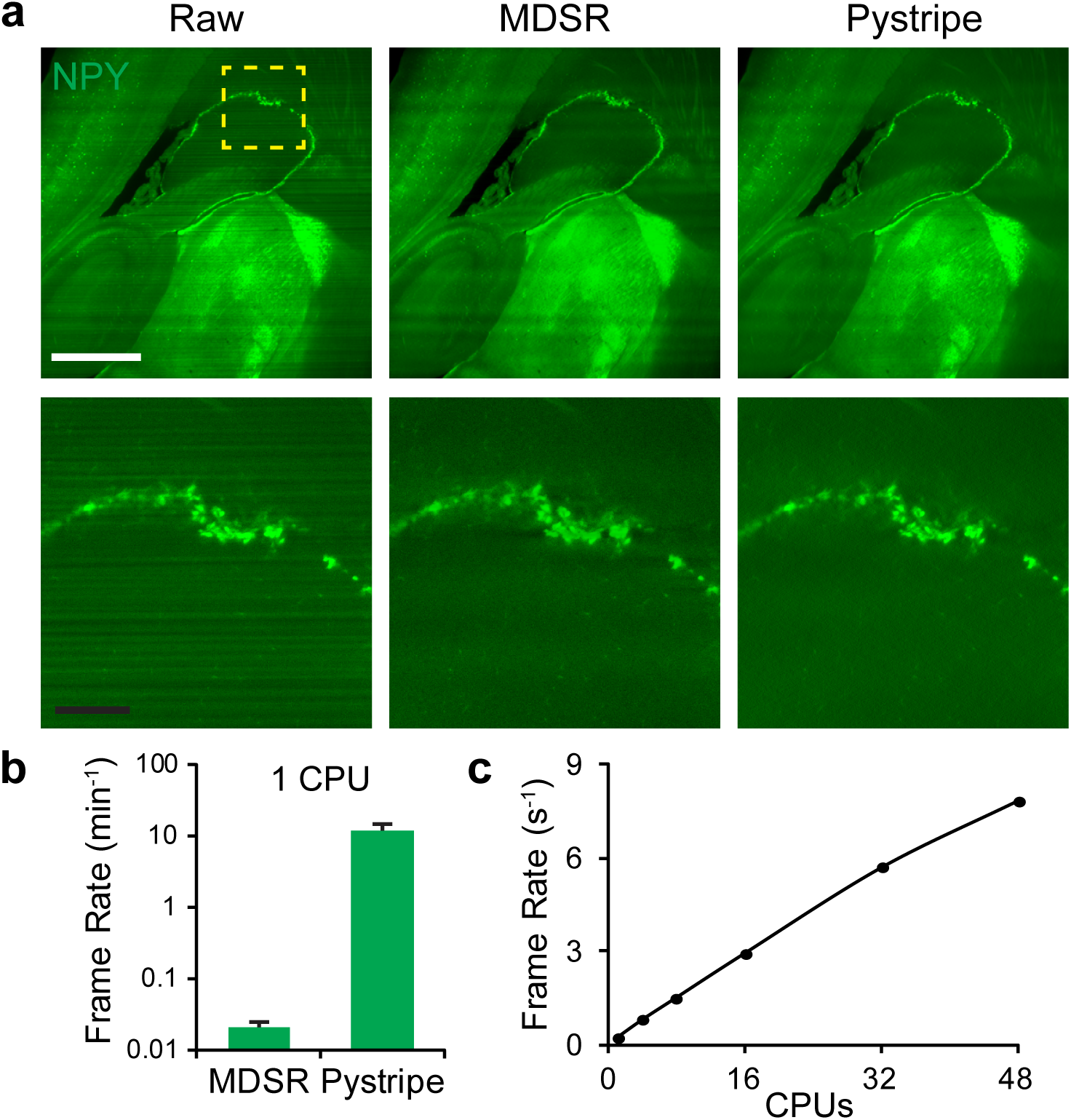
Destriping of light-sheet microscopy images using pystripe. (a) Comparison of destriping results from MDSR and pystripe on LSFM images of a SHIELD-processed mouse hemisphere stained for Neuropeptide Y (NPY). Scale bars, 1 mm (white) and 200 µm (black). (b) Average single-core destriping speed for MDSR and pystripe on 2048 x 2048 images (n = 10, error bars indicate standard deviation). (c) Scaling of destriping speed using pystripe with parallel processing on 2048 x 2048 images.

Pystripe also allows the user to provide an optional reference image for flat-field correction. Ideally, the reference flat would be calculated retroactively from the imaging data, but in practice a single flat for each channel of a particular imaging system still provides significant improvement. The example dataset includes a reference flat for illumination correction during the destriping step. By performing the illumination correction in pystripe, reading and writing the whole dataset multiple times can be avoided.

#### Stitching

Building on Terastitcher, TSV allows fast merging of stacks saved in Terastitcher hierarchal format. TSV obtains similar stitching quality as Terastitcher since it uses the same stack displacements and blending functions. Using TSV, a whole mouse hemisphere dataset was stitched with and without illumination correction and destriping using pystripe (Fig. 3a). Moderate vignetting effects were visible in the stitched original images at the intersections between adjacent stacks. These tiling artifacts were effectively reduced using flat-field correction in pystripe. Together, pystripe and TSV generate volumetric images that are ready for quantification by removing shadow and tiling artifacts before stitching (Fig. 3b, Supplementary Video 2). When comparing the stitching speed of TSV using multiple cores, the stitching frame rate increases linearly, reaching 2.6 frames per second with 48 cores (Fig. 3c).

**Figure 3.**
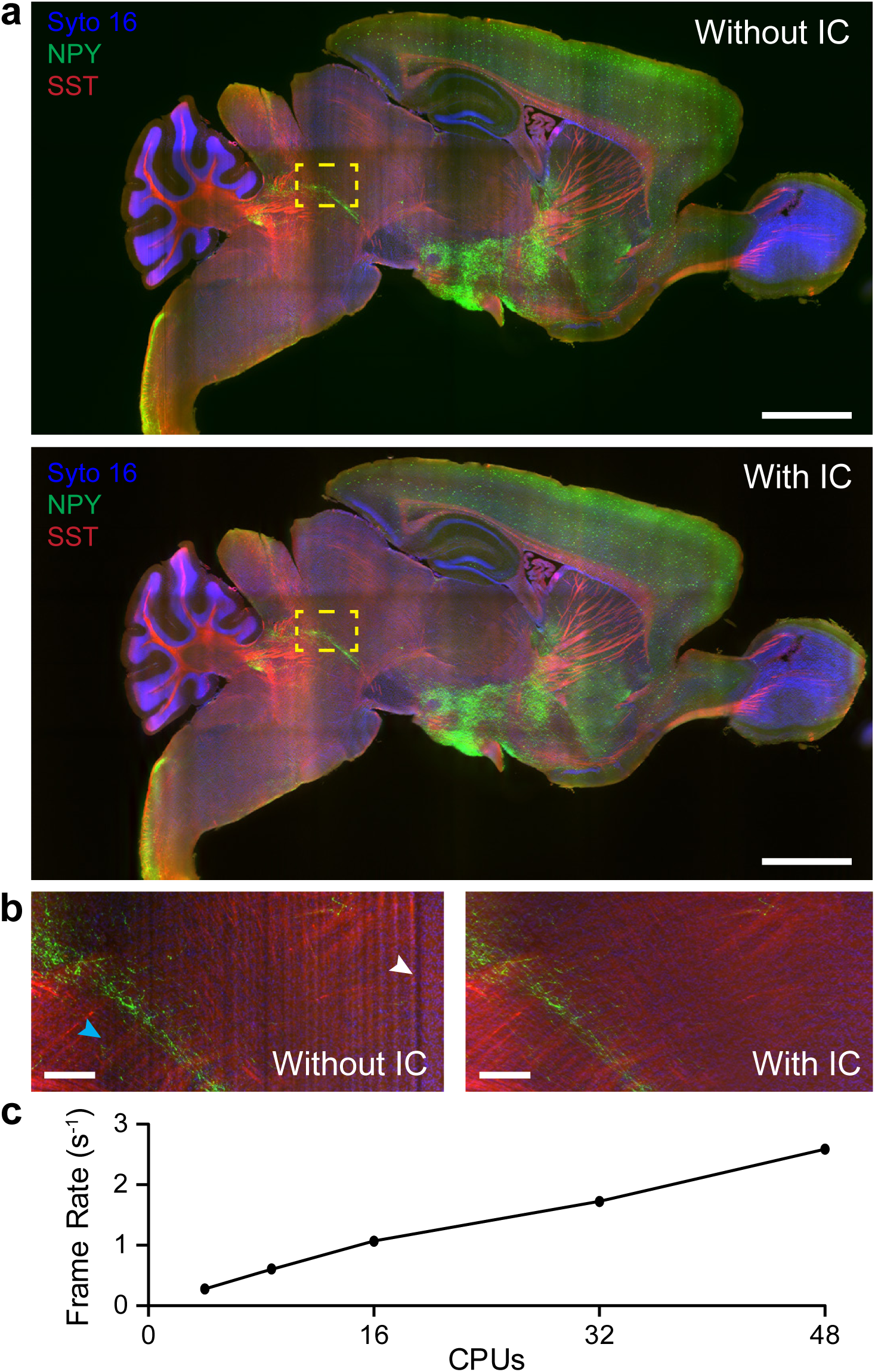
Stitching and illumination correction of light-sheet microscopy images using TSV. (a) Comparison of stitching results with and without destriping and illumination correction (IC) performed with pystripe on LSFM images of a SHIELD-processed mouse hemisphere. Scale bar, 2 mm. (b) Comparison of a region of interest with and without IC. Without IC, both uneven illumination (cyan arrow) and stripe artifacts (white arrow) corrupt the images. (c) Scaling of stitching speed using TSV with parallel processing on 2048 x 2048 images.

#### Atlas Alignment

Our hybrid automated atlas alignment method with manual refinement differs from wholly automated methods in that the alignment can be improved to the desired degree of accuracy via addition and modification of correspondences between the two volumes to be aligned (Fig. 4a). Tools for manual refinement are generally not used in combination with Elastix because integrating the transformations across multiple registration tools can be challenging. To the best of our knowledge, there are no web-based tools for interactive atlas alignment currently available.

**Figure 4.**
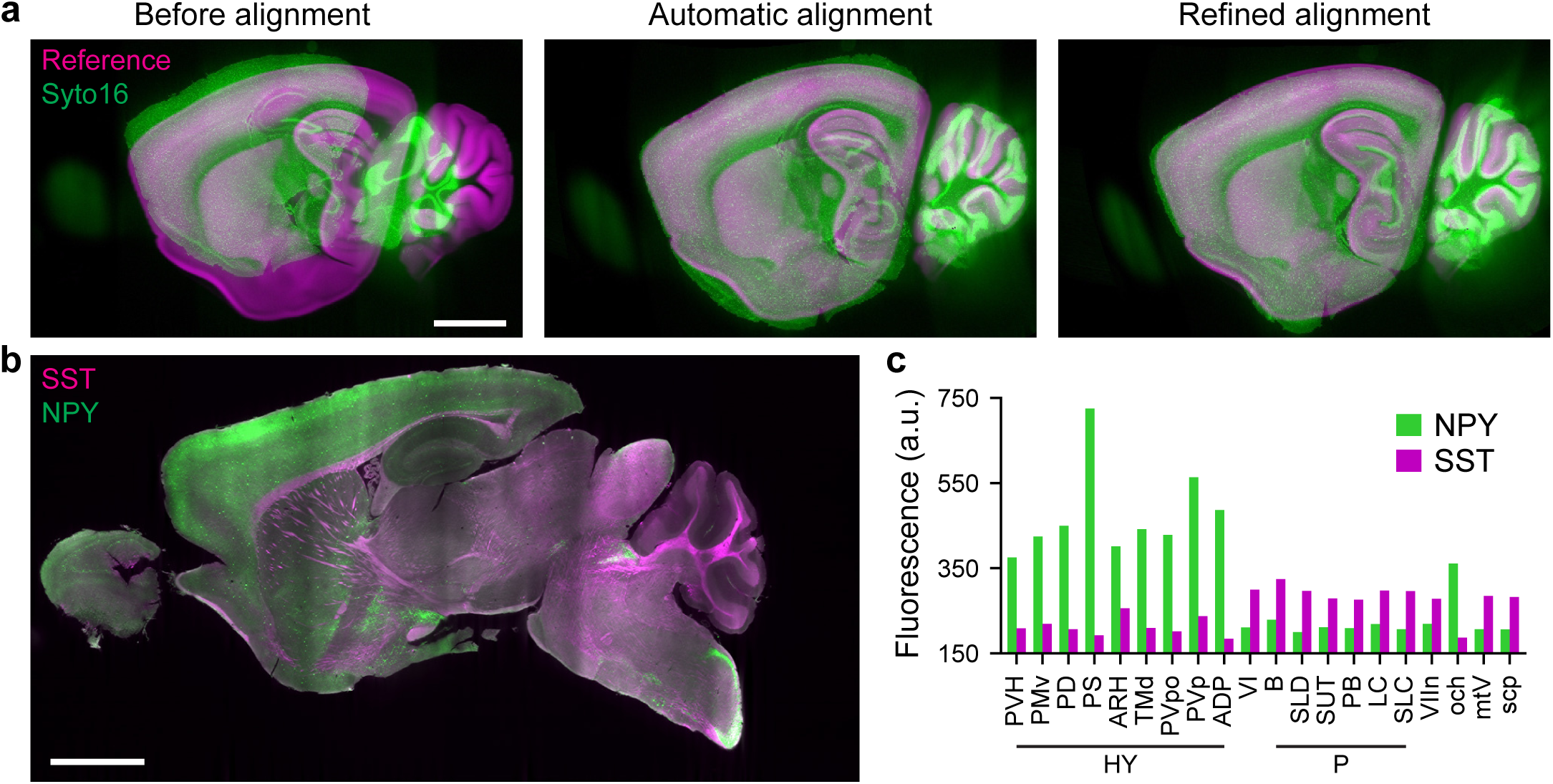
Atlas alignment and region-based fluorescence quantification using nuggt. (a) Comparison of atlas alignment of a syto 16 LSFM image to the reference image in the ABA before alignment, after automatic alignment, and after manual refinement. Scale bar, 2 mm. (b) Overlay of NPY and SST in the whole hemisphere example dataset (c) Mean fluorescence for the top 10 regions for both NPY and SST calculated after manual refinement. All other regions from the ABA are omitted for clarity.

After aligning a source autofluorescence or nuclear stain image to the reference image, other channels can be aligned to the atlas using the same calculated alignment. For example, neuropeptide Y (NPY) and somatostatin (SST) expression are included with syto 16 in separate channels of the provided example data (Fig. 4b). Using the alignment calculated by registering the syto 16 channel and the reference from the ABA, the mean fluorescence intensity of NPY and SST in each brain region can be calculated (Fig. 4c).

### Experimental design

All software modules are available from Github at http://www.github.com/chunglabmit/shield-2018 as well as from Docker hub at https://hub.docker.com/r/chunglabmit/shield-2018. We also provide example LSFM images of a SHIELD-processed mouse hemisphere dataset, which is available http://leviathan-chunglab.mit.edu/nature-protocols-2019. In order to adapt our image processing pipeline to other experimental situations, users should first complete our protocol using the provided example dataset. This dataset includes raw LSFM images as well as our intermediate results for users to compare and checkpoint their results throughout the pipeline. We also include a downsampled version of the full example dataset for users that would like to try our protocol on more modest computational hardware.

Young adult (2–4 months; median age 3 months) C57BL/6 mice were housed in a 12 h light/dark cycle with unrestricted access to food and water. To generate the example dataset, a single mouse brain was SHIELD-processed and stained with syto 16 and antibodies targeting NPY and SST using stochastic electrotransport^7^. The mouse brain sample was cut along the mid-sagittal plane and includes the olfactory bulb and the cerebellum. The stained hemisphere was then incubated in a RI-matching solution and imaged using a custom axially-swept LSFM system equipped with a 3.6x/0.2 objective (Special Optics). The resulting voxel width and depth are 1.8 µm and 2.0 µm, respectively. Although only one animal was involved in preparing the example dataset, our image processing protocol has been tested on over 15 intact mouse brain hemispheres from separate animals. All experimental protocols were approved by the MIT Institutional Animal Care and Use Committee and the Division of Comparative Medicine and were in accordance with guidelines from the National Institute of Health. All experiments using mice were conducted in strict adherence to the ethical regulations of MIT Institutional Animal Care and Use Committee and the Division of Comparative Medicine.

Our protocols have been developed for images of SHIELD-processed mouse brain tissues sectioned along the midsagittal plane with or without the olfactory bulb and cerebellum excised. The images must be acquired in a geometry that allows a transformation of axes (flipping, transposition) and cropping that bring the images into rough alignment with the atlas. In our experience, acquisition of either autofluorescence or a nuclear stain such as syto 16 provides enough mutual information for alignment with the reference volume for the ABA. Images were processed using only the techniques described in the protocol, and figures were prepared using linear lookup tables with adjustment of the minimum and maximum display range.

### Expertise needed to implement the protocol

Some minimal computer skills are needed to install Docker and navigate using the command line. If you are unfamiliar with Docker, important introductory information about Docker can be found at https://docs.docker.com/get-started/. Our protocol relies on several Docker features for volume sharing and network access, so a basic understanding of Docker is a prerequisite. Since Docker and most of our software is used from the command line, a basic understanding of how to use a terminal on the host operating system is also required.

### Limitations

Our overall image processing pipeline currently has been tested for analyzing mouse brain hemisphere datasets using the ABA autofluorescence reference volume. Therefore, we cannot guarantee that our overall pipeline will work in all cases for other species and atlases.

As previously mentioned, pystripe cannot remove striping artifacts that are not horizontally or vertically oriented within raw images. When processing images with very bright signals, the hybrid wavelet-FFT filter may introduce some ringing artifacts to the destriped images. The dual-band mode can mitigate these ringing artifacts, but in some cases, these artifacts may be undesirable. The user can elect to reduce the filter bandwidth or skip the destriping step in such cases.

When analyzing mouse brain hemispheres, severe tissue deformation due to improper sample preparation or handling may result in poor atlas alignment since the initial non-rigid alignment may not converge to a global optimum in such cases. Our atlas alignment protocol can be applied to mouse brain hemispheres with the olfactory bulb and cerebellum excised; however, this excision must be mirrored in the anatomical reference volume. This involves cropping the anatomical reference volume to match the source image, which is somewhat *ad hoc*. We provide reference volumes from the ABA for use in our Docker container along with reference volumes with the olfactory bulb, cerebellum, or both excised.

Processing whole-brain LSFM datasets is computationally expensive. We provide minimum system requirements that are recommended for processing our example mouse hemisphere dataset. However, we also provide a downsampled version of the original dataset for users without immediate access to computer hardware that meet the minimum system requirements.

## Materials

### EQUIPMENT

#### Example Data

All Example data is available from our laboratory servers (http://leviathan-chunglab.mit.edu/nature-protocols-2019), including:

- **Raw data**, comprising the raw LSFM images and a flat reference image from the provided mouse brain dataset.
- **Destriped data**, comprising the destriped LSFM images from the provided dataset.
- **Stitched data**, comprising the stitched destriped LSFM images from the provided dataset.
- **Alignment data**, containing the mean fluorescence in each region and intermediate results obtained during atlas alignment.
- **Downsampled data**, comprising our whole brain raw data downsampled and zipped for testing our pipeline on more modest computer hardware.

#### Computer Equipment

- For benchmarks, the computational pipeline was deployed on a workstation (TWS-1686525, Exxact) running Ubuntu 16.04 LTS on a 1 TB solid-state drive (Samsung) with two 24-core processors (Intel Xeon Platinum 8168), 768 GB of ECC memory, and a single 16 GB NVIDIA GPU (Tesla P100).
- Recommended minimum system requirements for processing the full-resolution example LSFM dataset. A computer with enough hard drive space to store the raw image data as well as the intermediate processing results is required. Our total example data is approximately 3 TB, suggesting that at least 4 TB of extra space is required. We recommend the following minimum system requirements in this case:
  - 8-core processor
  - 64 GB of memory
  - 256 GB solid-state drive with at least 32 GB available
  - 4 TB HDD available for data storage
- Recommended minimum system requirements for the processing the downsampled example LSFM dataset:
  - 2-core processor
  - 16 GB of memory
  - 128 GB solid-state drive with at least 32 GB available
- Software requirements.
  - Docker is recommended to run our software, but expert users can also directly install our software using the pip Python package manager from GitHub. If using our preconfigured Docker image to run our pipeline, Docker must be installed locally. Docker offers a free version of their software called Docker Community Edition.
  - We recommend using FIJI for inspecting images throughout our pipeline. FIJI can be obtained from https://fiji.sc/

### EQUIPMENT SETUP

#### Docker setup

1. Create a Docker ID to gain access to Docker software and Docker Hub using your preferred email address at https://hub.docker.com/. You may need to verify your email address.
2. Download Docker Community Edition for your particular operating system at https://www.docker.com/products/docker-engine.
3. Follow the installation instructions and the provided commands to verify that Docker has been installed correctly.
4. With the Docker deamon running, open a terminal and run the following command to install our preconfigured Docker image: docker pull chunglabmit/shield-2018 This command will download our software including all of the dependencies as well as commonly used resources needed for atlas alignment with the ABA. Note that we refer to the computer that is running Docker as the “host” and an instance of our Docker image as the “container”.

#### Downloading full resolution example data

This step requires 4 TB of available hard drive space if the whole dataset and intermediate results are downloaded. The raw data is approximately for each channel is approximately 560 GB. Proceed to *downloading downsampled example data* section if your computer does not have this much available space.

1. Create a folder named “data” to contain all of the example data on your machine and note its full path.
2. Start the Docker container with the data folder mounted by entering the following command into the command line: docker run -it -v path_to_data:/data chunglabmit/shield-2018 where “path_to_data” should be replaced with the full path to the data folder on the host. The command prompt should indicate that you are now the root user inside a running Docker container (as opposed to your usual username on the host). We refer to the command prompt inside a running container as the “Docker terminal window”.
3. Download the example data needed for a particular stage in the protocol by entering the following command(s) into the Docker terminal window. The data should begin to appear in the data folder that was created. Note that each download may take several hours.
  - Raw data (needed for destriping) wget -P /data -r --no-parent –Nh --cut-dirs 1 -R “index.html*” http://leviathan-chunglab.mit.edu/nature-protocols-2019/raw_data/
  - Destriped data (needed for stitching) wget -P /data -r --no-parent -nH --cut-dirs 1 -R “index.html*” http://leviathan-chunglab.mit.edu/nature-protocols-2019/destriped_data/
  - Stitched data (needed for atlas alignment) wget -P /data -r --no-parent -nH --cut-dirs 1 -R “index.html*” http://leviathan-chunglab.mit.edu/nature-protocols-2019/stitched_data/
  - Alignment data (for comparison of results) wget -P /data -r --no-parent -nH --cut-dirs 1 -R “index.html*” http://leviathan-chunglab.mit.edu/nature-protocols-2019/atlas/

#### Downloading downsampled example data

1. Navigate to http://leviathan-chunglab.mit.edu/nature-protocols-2019/ in your browser.
2. Click the downsampled_data.zip link and choose to save the file instead of opening it.
3. Unzip the downsampled_data.zip file to a new folder called “data” and make note of the full path of this folder on the host.

### Procedure

#### A) Destriping LSFM images using pystripe Timing 30 min setup (excluding download time), 1-4 h unattended computer time (depending on data size)

##### Container setup

i) Download the raw data (see ‘Downloading full resolution example data’ in the MATERIALS section) or move pre-existing raw data to a new folder called “data” and make note of the folder path on the host. Skip this step if using the downsampled example data. **CRITICAL STEP** Pystripe only accepts TIFF and RAW file formats. If using the full-resolution example data, the data folder should be created on a device with at least 4 TB of available space. **? TROUBLESHOOTING**
ii) Inspect the raw images using FIJI. **? TROUBLESHOOTING**
iii) Open a terminal and start the Docker container with the data folder from the host mounted inside the container using the following command: docker run -it -v path_to_data:/data chunglabmit/shield-2018 where “path_to_data” should be replaced with the full path of the data folder on the host. Note that the command prompt will change to the root user indicating that the prompt is now running interactively from within the container. **CRITICAL STEP** The semantics for mounting a volume to share data with the container are to specify a source path on the host and a target path inside the container. The syntax for expressing this at the command line is “-v path_on_host:path_in_container”. Note that path_in_container is a Unix-style path since the container is a Linux virtual machine. **CRITICAL STEP** Add quotes around the full path if it contains any spaces. **? TROUBLESHOOTING**
iv) Verify that the data folder is mounted correctly by running the following command from within the Docker terminal window. ls /data This command should list the contents of the shared data folder mounted in the container at the root path. **? TROUBLESHOOTING**

##### Destriping raw images from whole-brain LSFM

v) Run the following command to get the latest version of pystripe: git -C /shield-2018/pystripe pull Inspect the help message from pystripe by entering the following command from within the container. pystripe --help Instructions for using pystripe from the command line should print. Note from the instructions that an input folder and filter bandwidth are required and that pystripe will default to using all CPU cores available to the container.
vi) Destripe the raw data one channel at a time by entering the following commands within the Docker terminal window for each channel: pystripe -i /data/raw_data/channel -o /data/destriped_data/channel -s1 sigma1 -s2 sigma2 -w wavelet -c compression -x crossover -f flat.tif -d dark where “channel” represents the name of the folder containing images of the current channel, “sigma1” and “sigma2” represent the dual-band filter bandwidths in pixels (higher gives more filtering), “wavelet” is the mother wavelet name, “compression” is the amount of lossless TIFF compression, and “crossover” is the intensity range used to switch between filter bands. The arguments “flat” and “dark” are optional inputs for applying illumination correction. Including illumination correction in pystripe removes the need to load the images into memory multiple times. If using the full-resolution example data, then enter the following command. pystripe -i /data/raw_data/Ex_0_Em_0 -o /data/destriped_data/Ex_0_Em_0 -s1 256 -s2 256 -w db10 -c 3 -x 10 --flat flat.tif --dark 100 If using the downsampled example data, then enter the following command. pystripe -i /data/raw_data/Ex_0_Em_0 -o /data/destriped_data/Ex_0_Em_0 -s1 32 -s2 32 -w db10 -c 3 -x 10 --flat flat_downsampled.tif --dark 100 A progress bar showing the destriping progress for that channel will appear, and the destriped images will appear in the data folder, which is also available from the host machine. Inspect some of the resulting destriped images and the corresponding raw images in FIJI. Note that pystripe maintains the raw data folder structure in its output.
vii) Repeat steps A) vi-vii for all other channels.
viii) Shutdown the Docker container by typing “exit” in the Docker terminal window. The terminal will return to its original state before entering the Docker container. **CRITICAL STEP** Since Docker creates new containers each time they are started, no files inside the Docker container will be saved when restarting the container except for files in shared folders on the host, such as the mounted data folder. **PAUSE POINT** The destriped data folder can be archived for processing at a later date.

#### B) Stitching LSFM stacks using TSV TIMING 45 min setup (excluding download time), 4-12 h unattended computer time (depending on data size)

##### Container setup

i) Download the destriped data (see ‘Downloading full resolution example data’ in the MATERIALS section) or use the destriped data from the previous step. If using the downsampled data, complete part A and use the resulting destriped data for part B. **CRITICAL STEP** TSV only accepts TIFF and RAW file formats saved in Terastitcher hierarchical format. If using the full resolution example data, the data folder should be on a device with at least 4 TB of available space.
ii) Open a terminal and start the Docker container with the data folder from the host mounted inside the container using the following command: docker run -it -v path_to_data:/data chunglabmit/shield-2018 where “path_to_data” should be replaced with the full path of the data folder on the host. Note that the command prompt will change to the root user indicating that the prompt is now running interactively from within the container. **CRITICAL STEP** The semantics for mounting a volume to share data with the container are to specify a source path on the host and a target path inside the container. The syntax for expressing this at the command line is “-v path_on_host:path_in_container”. Note that path_in_container is a Unix-style path since the container is a Linux virtual machine. **CRITICAL STEP** Add quotes around the full path if it contains any spaces.

##### Stitching images from whole-brain LSFM

iii) Run the following command to get the latest version of TSV: git -C /shield-2018/tsv pull
iv) Use Terastitcher to create a stitching project file based on a single channel by entering the following command in the Docker terminal window. terastitcher −1 --volin=/data/destriped_data/channel_master --ref1=H --ref2=V --ref3=D --vxl1=x --vxl2=y --vxl3=z --projout=/data/destriped_data/channel_master/xml_import.xml --sparse_data where “channel_master” represents the name of the folder containing images of the channel used for calculating stack displacements and x, y, and z are the physical voxel dimensions in micron. If using the full-resolution example data, then enter the following command: terastitcher −1 --volin=/data/destriped_data/Ex_1_Em_1 --ref1=H --ref2=V --ref3=D --vxl1=1.8 --vxl2=1.8 --vxl3=2 --projout=/data/destriped_data/Ex_1_Em_1/xml_import.xml --sparse_data If using the downsampled example data, then enter the following command: terastitcher −1 --volin=/data/destriped_data/Ex_1_Em_1 --ref1=H --ref2=V --ref3=D --vxl1=14.4 --vxl2=14.4 --vxl3=16 --projout=/data/destriped_data/Ex_1_Em_1/xml_import.xml --sparse_data **CRITICAL STEP** The voxel dimensions must match the voxel dimensions for the LSFM system in microns. **? TROUBLESHOOTING** v) Calculate the stack displacements using Terastitcher by entering the following command in the Docker terminal window: terastitcher −2 --projin=/data/destriped_data/channel_master/xml_import.xml --projout=/data/destriped_data/channel_master/xml_displacement.xml If using the example data, enter the following command: terastitcher −2 --projin=/data/destriped_data/Ex_1_Em_1/xml_import.xml --projout=/data/destriped_data/Ex_1_Em_1/xml_displacement.xml **? TROUBLESHOOTING**
vi) Generate a Terastitcher project file by entering the following command in the Docker terminal window: terastitcher −3 --projin=/data/destriped_data/channel_master/xml_displacement.xml --projout=/data/destriped_data/channel_master/xml_displproj.xml If using the example data, enter the following command: terastitcher −3 --projin=/data/destriped_data/Ex_1_Em_1/xml_displacement.xml --projout=/data/destriped_data/Ex_1_Em_1/xml_displproj.xml
vii) Use TSV to generate stitched images by entering the following command in the Docker terminal window: tsv-convert-2D-tif --xml-path /data/destriped_data/channel_master/xml_displproj.xml --output-pattern /data/stitched_data/channel_master/”{z:04d}.tiff” --compression compression --ignore-z-offsets where “/data/stitched_data/channel_master” is a new folder that will be created for the stitched images from the master channel, and “compression” is the amount of lossless TIFF compression to use. If using the example data, enter the following command: tsv-convert-2D-tif --xml-path /data/destriped_data/Ex_1_Em_1/xml_displproj.xml --output-pattern /data/stitched_data/Ex_1_Em_1_master/”{z:04d}.tiff” --compression 4 --ignore-z-offsets A progress bar will appear showing the stitching progress. Note that TSV will default to using all processor cores available to the container.
viii) Inspect the stitched images in FIJI. TSV generates TIFF images which can be imported directly into FIJI.
ix) If there are no more channels to stitched, then proceed to part C of the protocol. If there are more channels to be stitched, then the following command will stitch other channels using the displacements calculated from the master channel: tsv-convert-2D-tif --xml-path /data/destriped_data/channel_master/xml_displproj.xml --output-pattern /data/stitched_data/channel/”{z:04d}.tiff” --compression compression --ignore-z-offsets --input /data/destriped_data/channel Where “channel_master” represents the folder name of the channel used for calculating stack displacements, “channel” represents folder of images from another channel to be stitched using the previously computed displacements, and the other arguments are as described before. Repeat this command for all other channels to be stitched. If using the example data, then enter the following commands: tsv-convert-2D-tif --xml-path /data/destriped_data/Ex_1_Em_1/xml_displproj.xml --output-pattern /data/stitched_data/Ex_0_Em_0/“{z:04d}.tiff” --compression 4 --ignore-z-offsets --input /data/destriped_data/Ex_0_Em_0 tsv-convert-2D-tif --xml-path /data/destriped_data/Ex_1_Em_1/xml_displproj.xml --output-pattern /data/stitched_data/Ex_2_Em_2/“{z:04d}.tiff” --compression 4 --ignore-z-offsets --input /data/destriped_data/Ex_2_Em_2
x) Close the Docker container by typing “exit” into the Docker terminal window and pressing “Enter”. **PAUSE POINT** The stitched data folder can be archived for processing at a later date.

#### c) Atlas alignment with manual refinement using nuggt TIMING 1-2 hr setup (excluding download time), 6-12 h unattended computer time (depending on data set size)

##### Container setup and Neuroglancer basics

i) Download the stitched data (see ‘Downloading full resolution example data’ in the MATERIALS section) or use the stitched data from the previous step. If using the downsampled data, complete part B and use the resulting stitched data for part C.
ii) For Linux users, open a terminal and start the Docker container with the data folder from the host mounted inside the container using the following command: docker run -it --expose 8999 -p 8999:8999/tcp --network host -v path_to_data:/data --shm-size 1g chunglabmit/shield-2018 For Windows and Mac users, enter the following command: docker run -it --expose 8999 -p 8999:8999/tcp -v path_to_data:/data --shm-size 1g chunglabmit/shield-2018 **CRITICAL STEP** Add quotes around the full path if it contains any spaces.
iii) The following files are included in the “/allen-mouse-brain-atlas” folder inside the Docker container:
  - **autofluorescence_25_half_sagittal.tif** -a 3D image of the autofluorescence channel for the mid-sagittal sectioning of the Allen Mouse Brain atlas reference (to be used in place of “/reference/reference.tiff”).
  - **annotation_25_half_sagittal.tif** -a 3D segmentation of the Allen Mouse Brain reference (to be used in place of “/reference/segmentation.tiff”).
  - **coords_25_half_sagittal.json** -the coordinates of key points on the reference (to be used in place of “/reference/points.json”)
  - **AllBrainRegions.csv** -a mapping of region ID numbers in the segmentation to names of regions.
  Use nuggt to inspect the included mouse brain reference atlas by entering the following command into the Docker terminal window: nuggt-display --port 8999 [--ip-address 0.0.0.0] --segmentation /allen-mouse-brain-atlas/annotation_25_half_sagittal.tif --points /allen-mouse-brain-atlas/coords_25_half_sagittal.json /allen-mouse-brain-atlas/autofluorescence_25_half_sagittal.tif reference gray where “--ip-address 0.0.0.0” should be included by Windows and Mac users. The nuggt-display application will print a URL similar to “http://127.0.0.1:8999/v/…” for Linux users in the Docker terminal window. For Windows and Mac users, the nuggt-display application will print a URL similar to ” http://<container-id>:8999/v/…” in the Docker terminal window, and the container ID should be replaced with “localhost” in the next step.
iv) Open the URL in your browser to see the Neuroglancer user interface. The browser should display four panels. The top left one is the XY view, the top right is the XZ view, the bottom right one is the YZ view, and the bottom left one is a 3D view. The current cursor position in image coordinates is displayed at the top of the Neuroglancer user interface. A list of keyboard shortcuts is available if you type “h” while using Neuroglancer. **? TROUBLESHOOTING** In Neuroglancer, you can move the stack by pressing the left mouse button with the cursor in any of the three orthogonal view panels and dragging. You can scroll through the depth of the stack by turning the mouse wheel, and you can zoom in and out by pressing the “ctrl” key and turning the mouse wheel. Each layer in Neuroglancer listed along the top of the user interface can be hidden by clicking the layer name. Segmentation layers can used to highlight specific brain regions by double-clicking regions in any view panel.
v) Exit the nuggt-display application by holding “ctrl” and pressing “C” in the Docker terminal window.

##### Create a points file for a custom reference atlas

This step is optional if you are aligning a sagittal image of a mouse brain hemisphere to the Allen Mouse Brain Atlas (which is the case when using the example data). If you are using the example data, proceed to step C xii.

vi) Locate either a 3D TIFF file of the reference image (e.g. “path_to_reference/reference.tiff”) or a folder of 2D TIFF files named in strict ascending alphabetical order (e.g. “path_to_reference/images/img_0000.tiff”) on the host. **CRITICAL STEP** You must prepare an accompanying segmentation of this reference (e.g. “path_to_reference/segmentation.tiff”) where each pixel of the segmentation has a value corresponding to the region in which the pixel lies. You must also have a mapping of region number to the region’s name in a format similar to the file “/allen-mouse-brain-atlas/AllBrainRegions.csv”.
vii) Open a terminal and start the Docker container with the reference image(s) mounted using the following command: docker run -it --expose 8999 -p 8999:8999/tcp [--network host] -v path_to_reference:/reference -v path_to_data:/data chunglabmit/shield-2018 where “path_to_reference” and “path_to_data” represent the full paths of the directories containing the reference image(s) and the images to be aligned to the reference on the host. The “--network host” argument should only be included by Linux users. **CRITICAL STEP** Add quotes around the full path if it contains any spaces.
viii) If you have a 3D reference TIFF file, type the following command into the Docker terminal window: nuggt --port 8999 --image /reference/reference.tiff --output /reference/points.json where “reference.tiff” should be replaced with the name of your reference image. If you have a folder of 2D TIFF files, type: nuggt --port 8999 --image “/reference/images/*.tiff” --output /reference/points.json **CRITICAL STEP** Use “*.tif” instead of “*.tiff” if that is the extension of your image files.
ix) The nuggt application will display a line similar to “Editing viewer: http://127.0.0.1:8999/v/…” in the Docker terminal window. Windows and Mac users should replace the container ID in the URL with “localhost”. Open the URL in your browser to see the Neuroglancer user interface.
x) Add fiducial points by placing the cursor where you want the point to be, holding the “shift” key and pressing “A”. You can delete a point by placing the cursor over it, holding the shift key and pressing “D”. You should annotate the image using easily identifiable locations like the dentate gyrus as well as the perimeter of the image (see Supplementary Fig. 1). **? TROUBLESHOOTING**
xi) Hold the “shift” key and press “S” to save the annotations. Bring up the Docker terminal window again, hold “ctrl” and press “C” to exit from the nuggt application. **PAUSE POINT** The points.json file can be saved and reused for future atlas alignment tasks using the same reference image(s).

##### Perform automatic alignment

xii) Determine whether the images to be aligned with the reference atlas needs to be flipped or rotated by inspecting the stitched and reference images. If aligning to the provided sagittal mouse brain atlas, each z-plane of the stitched images should be approximately a sagittal section with the olfactory bulb at the top of the image and the cortex to the right (see Supplementary Fig. 1). The z-plane corresponding to the medial sagittal section should be the last image in the stack (the one that is the last in alphabetical order if 2D planes are used). If the cortex is at the left, you will need to “flip-X”. If the olfactory bulb is at the bottom, you will need to “flip-Y”. If the medial sagittal section is at Z = 0, then you will need to “flip-Z”. Note that “flipping” in this case means reversing the index of an image dimension, which corresponds to a reflection of the image in that dimension. **? TROUBLESHOOTING** If aligning the example data to the provided mouse brain atlas, then the Ex_0_Em_0 channel (containing syto 16 nuclear stain) should be used for atlas alignment and flipped in the X and Z dimensions (see Supplementary Fig. 2).
xiii) Determine whether the channel to be aligned with the reference atlas needs to be cropped. Typically, the reference image extends to the left, right, top and bottom with no background margins, but the stitched images to be aligned have a margin to the left, right, top or bottom. If these margins are large, they will prevent the automatic alignment from succeeding; thus, the image to be aligned must be cropped to match when performing the automated alignment. Note the X-start and X-end, Y-start and Y-end, and Z-start and Z-end (if whole planes need to be cropped) coordinates of the image after cropping using FIJI to examine the images. **CRITICAL STEP** These cropping coordinates apply to the image before flipping. If using the full resolution example data, then use the following crop coordinates:
  - X-start, X-stop: None
  - Y-start, Y-stop: 0, 9800
  - Z-start, Z-stop: None
  If using the downsampled example data, then use the following crop coordinates:
  - X-start, X-stop: None
  - Y-start, Y-stop: 0, 1225
  - Z-start, Z-stop: None
xiv) Run the rescale-image-for-alignment program to rescale, flip, and crop the channel used for atlas alignment. If step (xiii) determined that the image needs X-flipping, use the --flip-x switch, and similarly for Y and Z. Likewise, if step (xiii) determined the image needs X-cropping, use the --clip-x argument, and similarly for Y and Z. For example, if an image needs to be flipped in X and Z and cropped in Y at the bottom of the image at coordinate 9100, the resulting command to be typed into the Docker terminal window is: rescale-image-for-alignment --input “/data/stitched_data/channel_alignment/*.tiff” --atlas-file /reference/reference.tiff --output /data/downsampled_flip-x_flip-z_clip-y-0-9100.tiff --flip-x --flip-z --clip-y 0,9100 where “channel_alignment” is the folder containing the stitched images of the channel used for alignment. If using the full resolution example data with the provided reference atlas, then type: rescale-image-for-alignment --input “/data/stitched_data/Ex_0_Em_0/*.tiff” --atlas-file /allen-mouse-brain-atlas/autofluorescence_25_half_sagittal.tif –output /data/downsampled_flip-x_flip-z_clip-y-0-9800.tiff --flip-x --flip-z --clip-y 0,9800 If using the downsampled example data with the provided reference atlas, then type: rescale-image-for-alignment --input “/data/stitched_data/Ex_0_Em_0/*.tiff” --atlas-file /allen-mouse-brain-atlas/autofluorescence_25_half_sagittal.tif –output /data/downsampled_flip-x_flip-z_clip-y-0-1225.tiff --flip-x --flip-z --clip-y 0,1225 **CRITICAL STEP** Record the values for the --flip-x, --flip-y, --flip-z --clip-x, --clip-y and --clip-z switches in the output image name. These parameters will be used again in step C xxiv. **CRITICAL STEP** If using the example data, make sure that Ex_0_Em_0, the syto 16 channel, is rescaled and used for atlas alignment.
xv) Align the rescaled image to the reference using Elastix. If using a custom reference image in the same example scenario presented in step C xiv, then the following command would be used: sitk-align --moving-file /data/downsampled_flip-x_flip-z_clip-y-0-9100.tiff --fixed-file /reference/reference.tiff --fixed-point-file /reference/points.json --xyz --alignment-point-file /data/alignment.json **CRITICAL STEP** Substitute the name of your reference image for “reference.tiff” above and substitute the name you used as the --moving-file in step (C)(xiii) for “downsampled.tiff” above. If using the full resolution example data with the provided reference atlas, then type: sitk-align --moving-file /data/downsampled_flip-x_flip-z_clip-y-0-9800.tiff --fixed-file /allen-mouse-brain-atlas/autofluorescence_25_half_sagittal.tif --fixed-point-file /allen-mouse-brain-atlas/coords_25_half_sagittal.json --xyz --alignment-point-file /data/alignment.json If using the downsampled example data with the provided reference atlas, then type: sitk-align --moving-file /data/downsampled_flip-x_flip-z_clip-y-0-1225.tiff --fixed-file /allen-mouse-brain-atlas/autofluorescence_25_half_sagittal.tif --fixed-point-file /allen-mouse-brain-atlas/coords_25_half_sagittal.json --xyz --alignment-point-file /data/alignment.json **? TROUBLESHOOTING**

##### Refine the alignment manually

xvi) Use nuggt-align to manually refine the automatic alignment. If using a custom reference image in the same example scenario presented in step C xiv, then the following command would be used: nuggt-align --port 8999 [--ip-address 0.0.0.0] --reference-image /reference/reference.tiff --moving-image /data/downsampled_flip-x_flip-z_clip-y-0-9100.tiff –points /data/alignment.json where “--ip-address 0.0.0.0” should be included by Windows and Mac users. **CRITICAL STEP** Substitute the name of your reference image for “reference.tiff” above and substitute the name you used as the --moving-file in step (C)(xiii) for “downsampled.tiff” above. If aligning the full resolution example data to the provided reference atlas, then enter the following command in the Docker terminal window: nuggt-align --port 8999 [--ip-address 0.0.0.0] --reference-image /allen-mouse-brain-atlas/autofluorescence_25_half_sagittal.tif --moving-image downsampled_flip-x_flip-z_clip-y-0-9800.tiff --points /data/alignment.json If aligning the downsampled example data to the provided reference atlas, then enter the following command in the Docker terminal window: nuggt-align --port 8999 [--ip-address 0.0.0.0] --reference-image /allen-mouse-brain-atlas/autofluorescence_25_half_sagittal.tif --moving-image /data/downsampled_flip-x_flip-z_clip-y-0-1225.tiff --points /data/alignment.json
xvii) There are two links that are displayed by nuggt-align in the Docker terminal window. One is the link to a Neuroglancer webpage displaying the reference image and one is a link to the image that is to be aligned. Copy each of the links and open them in separate windows in your web browser. Windows and Mac users should change the container ID in the URL to “localhost” (see step C iii).
xviii) Refine the locations of each of the fiducial points in the image. This is done by selecting the reference image browser window (e.g. by clicking in it), holding down the “shift” key and typing “N”. A red point will appear in both the reference and moving image browsers. You should adjust the point in the moving image browser to correspond with its location in the reference image if it is placed incorrectly. This is done by holding down the “ctrl” key and clicking on the location in the reference image where the point should be. The point can be readjusted multiple times until you are satisfied with the location. After this is done, hold the shift key down and type, “D” (for “done”) and hold the shift key down and type “N” to move to the next fiducial point. Repeat the adjustment procedure until all of the fiducial points have been adjusted. **? TROUBLESHOOTING**
xix) Hold the shift key down and press “S” to save the adjusted alignment. **CRITICAL STEP** The location of the refined fiducial points are not saved automatically. The refined atlas alignment progress will be lost if the adjusted alignment points are not saved.
xx) Hold the shift key down and type “W” to warp the image to be aligned to the reference image. **CRITICAL STEP** Wait for the message, “Warping complete”, to be displayed before doing anything further.
xxi) Add additional points to refine the warping further. Add a point to the reference image by holding the “ctrl” key down and pressing the mouse button while hovering over the location of a discrepancy. Focus the alignment image to this location by holding down the “shift” key and pressing “T” (for “translate”). In the alignment image Neuroglancer webpage, adjust the location of the point by holding down the “ctrl” key and pressing the left mouse button, then hold the shift key down and press “D” (for “Done”). **CRITICAL STEP** Adding duplicate points in the same location will cause the warping to fail. To avoid this problem, it is helpful to warp the moving image after adding each point to verify that the warping is successful. **? TROUBLESHOOTING**
xxii) Hold the shift key down and press “S” to save the adjusted alignment. **CRITICAL STEP** The location of the refined fiducial points are not saved automatically. The refined atlas alignment progress will be lost if the adjusted alignment points are not saved.
xxiii) In the Docker terminal window, hold down the “ctrl” key and press “C” to exit the nuggt-align program. xxiv) Rescale the alignment to the size of your original image. If using a custom reference image in the same example scenario presented in step C xiv, then the following command would be used: rescale-alignment-file --stack “/data/stitched_data/channel_alignment/*.tiff” --alignment-image /data/downsampled_flip-x_flip-z_clip-y-0-9100.tiff --input /data/alignment.json --output /data/rescaled-alignment.json --flip-x --flip-z --clip-y 0,9100 **CRITICAL STEP** replace “--flip-x --flip-z --clip-y 0,9100” above with the values used in step C xiv If aligning the full resolution example data to the provided reference atlas, then enter the following command in the Docker terminal window: rescale-alignment-file --stack “/data/stitched_data/Ex_0_Em_0/*.tiff” --alignment-image /data/downsampled_flip-x_flip-z_clip-y-0-9800.tiff --input /data/alignment.json –output /data/rescaled-alignment.json --flip-x --flip-z --clip-y 0,9800 If aligning the downsampled example data to the provided reference atlas, then enter the following command in the Docker terminal window: rescale-alignment-file --stack “/data/stitched_data/Ex_0_Em_0/*.tiff” --alignment-image /data/downsampled_flip-x_flip-z_clip-y-0-1225.tiff --input /data/alignment.json –output /data/rescaled-alignment.json --flip-x --flip-z --clip-y 0,1225 **PAUSE POINT** The alignment.json and rescaled-alignment.json files can be saved for continuing the analysis at a later date.

##### Compute the total and mean intensity per region

xxv) Use the rescaled alignment file to compute fluorescence statistics of each channel for all brain regions. If using a custom reference image, then enter the following command into the Docker terminal window: calculate-intensity-in-regions --input “/data/stitched_data/channel/*.tiff” --alignment /data/rescaled-alignment.json --reference-segmentation /reference/segmentation.tiff --brain-regions-csv /reference/brain-regions.csv --output /data/results-level-7.csv --level 7 where “channel” represents the folder of the current channel to be analyzed. **CRITICAL STEP** the “--level” argument picks the granularity of the segmentation description from 7 (finest granularity) to 1 (coarsest granularity). You can specify multiple levels to be calculated at the same time at little additional computational cost. If aligning the full resolution example data to the provided reference atlas, then enter the following command: calculate-intensity-in-regions --input “/data/stitched_data/Ex_1_Em_1_master/*.tiff” --alignment /data/rescaled-alignment.json --reference-segmentation /allen-mouse-brain-atlas/annotation_25_half_sagittal.tif --brain-regions-csv /allen-mouse-brain-atlas/AllBrainRegions.csv --output /data/results-level-3.csv --level 3 –output /data/results-level-4.csv --level 4 --output /data/results-level-5.csv --level 5 –output /data/results-level-6.csv --level 6 --output /data/results-level-7.csv --level 7 --shrink 2 –n-cores N where N is the number of cores to use for calculating fluorescence statistics. Note that using more cores causes the memory usage to increase as well. If memory becomes a limiting factor, then limit the number of cores used or increase the shrink factor if acceptable. If aligning the downsampled example data to the provided reference atlas, then enter the following command: calculate-intensity-in-regions --input “/data/stitched_data/Ex_1_Em_1_master/*.tiff” --alignment /data/rescaled-alignment.json --reference-segmentation /allen-mouse-brain-atlas/annotation_25_half_sagittal.tif --brain-regions-csv /allen-mouse-brain-atlas/AllBrainRegions.csv --output /data/results-level-3.csv --level 3 –output /data/results-level-4.csv --level 4 --output /data/results-level-5.csv --level 5 –output /data/results-level-6.csv --level 6 --output /data/results-level-7.csv --level 7 This command can be repeated for all other channels to be analyzed. The results file that is created contains a spreadsheet of the region names, the volume of each region in voxels, the sum of all intensities in each region (total_intensity) and the average intensity of the voxels in each region (mean_intensity).

### Timing

For processing the full resolution example data:

Stage A, 30 min for setup and 2-4 h of unattended computer time

Stage B, 45 min for setup and 4-12 h of unattended computer time

Stage C, 1-2 h for setup and manual refinement and 4-12 h of unattended computer time For processing the downsampled example data:

Stage A, 30 min for setup and 10 min of unattended computer time

Stage B, 45 min for setup and 20 min of unattended computer time

Stage C, 1-2 h for setup and manual refinement and 30 min of unattended computer time

### Troubleshooting

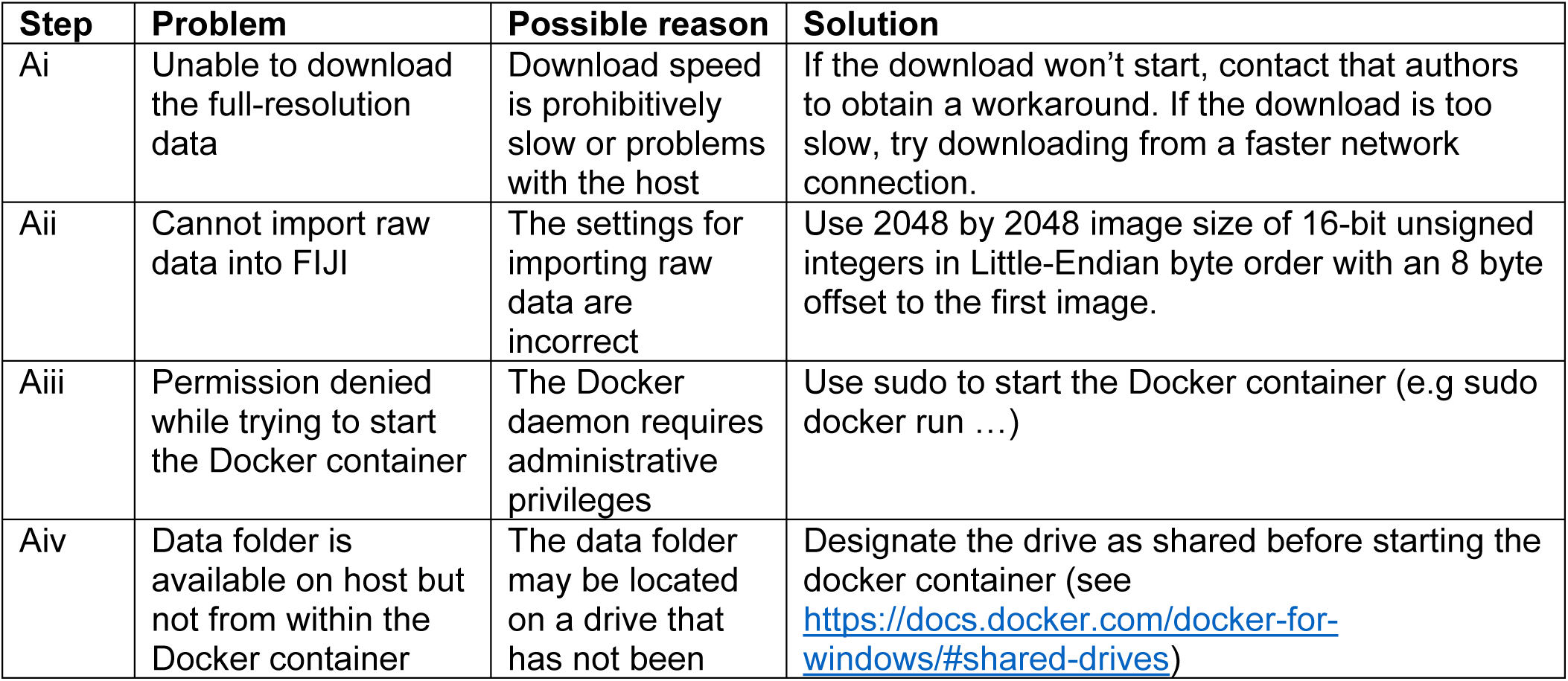
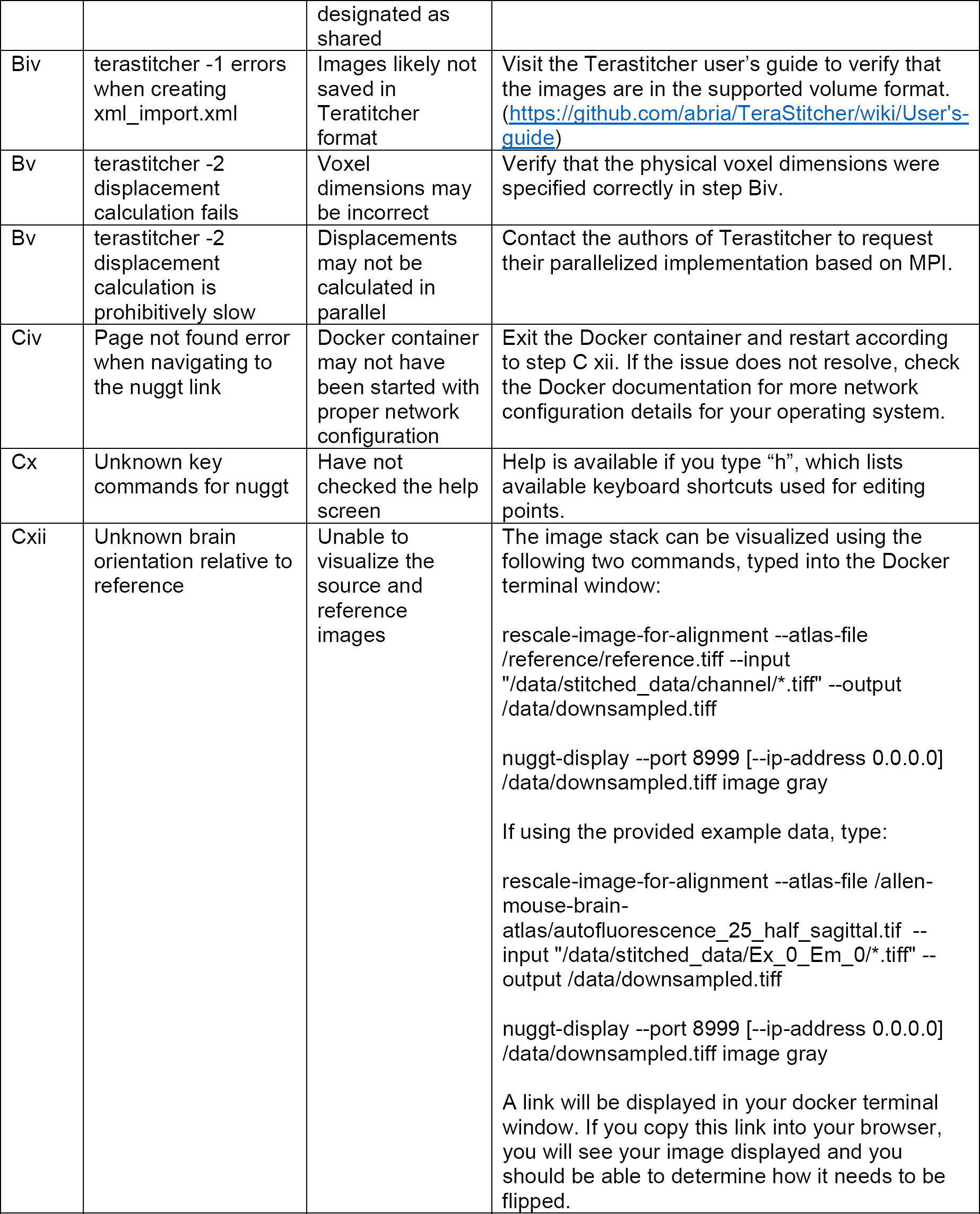
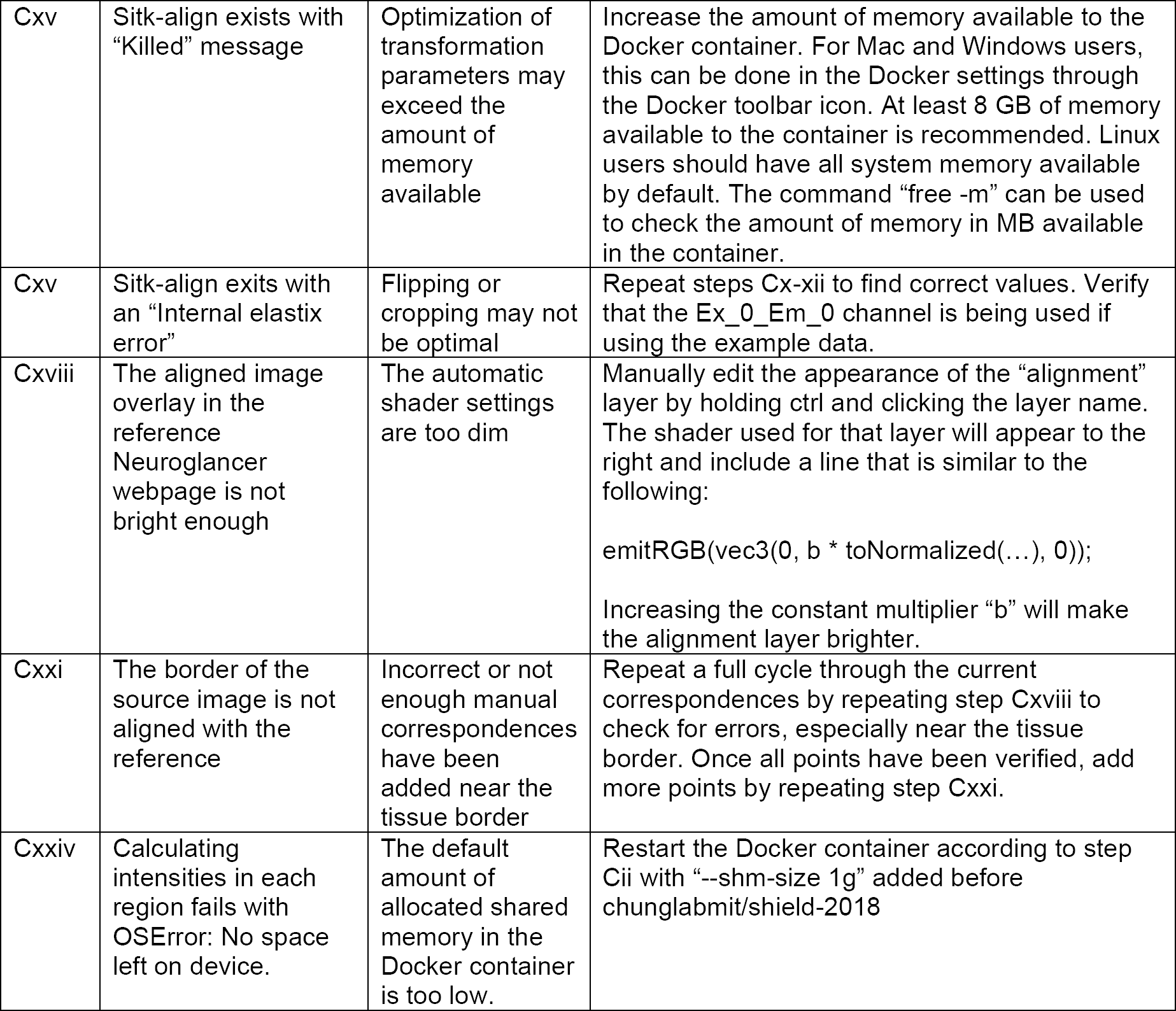

| Step | Problem | Possible reason | Solution |
| --- | --- | --- | --- |
| Ai | Unable to download the full-resolution data | Download speed is prohibitively slow or problems with the host | If the download won't start, contact that authors to obtain a workaround. If the download is too slow, try downloading from a faster network connection. |
| Aii | Cannot import raw data into FIJI | The settings for importing raw data are incorrect | Use 2048 by 2048 image size of 16-bit unsigned integers in Little-Endian byte order with an 8 byte offset to the first image. |
| Aiii | Permission denied while trying to start the Docker container | The Docker daemon requires administrative privileges | Use sudo to start the Docker container (e.g sudo docker run ...) |
| Aiv | Data folder is available on host but not from within the Docker container | The data folder may be located on a drive that has not been designated as shared | Designate the drive as shared before starting the docker container (see https://docs.docker.com/docker-for-windows/#shared-drives) |
| Biv | terastitcher -1 errors when creating xml_import.xml | Images likely not saved in Teratitcher format | Visit the Terastitcher user's guide to verify that the images are in the supported volume format. (https://github.com/abria/TeraStitcher/wiki/User's-guide) |
| Bv | terastitcher -2 displacement calculation fails | Voxel dimensions may be incorrect | Verify that the physical voxel dimensions were specified correctly in step Biv. |
| Bv | terastitcher -2 displacement calculation is prohibitively slow | Displacements may not be calculated in parallel | Contact the authors of Terastitcher to request their parallelized implementation based on MPI. |
| Civ | Page not found error when navigating to the nuggt link | Docker container may not have been started with proper network configuration | Exit the Docker container and restart according to step C xii. If the issue does not resolve, check the Docker documentation for more network configuration details for your operating system. |
| Cx | Unknown key commands for nuggt | Have not checked the help screen | Help is available if you type "h", which lists available keyboard shortcuts used for editing points. |
| Cxii | Unknown brain orientation relative to reference | Unable to visualize the source and reference images | The image stack can be visualized using the following two commands, typed into the Docker terminal window:<br>`rescale-image-for-alignment --atlas-file /reference/reference.tiff --input "/data/stitched_data/channel/*.tiff" --output /data/downsampled.tiff` `nuggt-display --port 8999 [--ip-address 0.0.0.0] /data/downsampled.tiff image gray`<br>If using the provided example data, type:<br>`rescale-image-for-alignment --atlas-file /allen-mouse-brain-atlas/autofluorescence_25_half_sagittal.tif --input "/data/stitched_data/Ex_0_Em_0/*.tiff" --output /data/downsampled.tiff` `nuggt-display --port 8999 [--ip-address 0.0.0.0] /data/downsampled.tiff image gray`<br>A link will be displayed in your docker terminal window. If you copy this link into your browser, you will see your image displayed and you should be able to determine how it needs to be flipped. |
| Cxv | Sitk-align exists with “Killed” message | Optimization of transformation parameters may exceed the amount of memory available | Increase the amount of memory available to the Docker container. For Mac and Windows users, this can be done in the Docker settings through the Docker toolbar icon. At least 8 GB of memory available to the container is recommended. Linux users should have all system memory available by default. The command “free -m” can be used to check the amount of memory in MB available in the container. |
| Cxv | Sitk-align exits with an “Internal elastix error” | Flipping or cropping may not be optimal | Repeat steps Cx-xii to find correct values. Verify that the Ex_0_Em_0 channel is being used if using the example data. |
| Cxviii | The aligned image overlay in the reference Neuroglancer webpage is not bright enough | The automatic shader settings are too dim | Manually edit the appearance of the “alignment” layer by holding ctrl and clicking the layer name. The shader used for that layer will appear to the right and include a line that is similar to the following:<br>emitRGB(vec3(0, b * toNormalized(...), 0));<br>Increasing the constant multiplier “b” will make the alignment layer brighter. |
| Cxxi | The border of the source image is not aligned with the reference | Incorrect or not enough manual correspondences have been added near the tissue border | Repeat a full cycle through the current correspondences by repeating step Cxviii to check for errors, especially near the tissue border. Once all points have been verified, add more points by repeating step Cxxi. |
| Cxxiv | Calculating intensities in each region fails with OSError: No space left on device. | The default amount of allocated shared memory in the Docker container is too low. | Restart the Docker container according to step Cii with “--shm-size 1g” added before chunglabmit/shield-2018 |

#### Anticipated Results

Upon successful completion of our image processing pipeline, our protocol will yield corrected and stitched multichannel volumetric images of a whole mouse brain hemisphere as well as an alignment with a provided atlas for brain region segmentation. The atlas alignment is used to create spreadsheets in CSV format containing the volume, total fluorescence, and mean fluorescence of each brain region and each channel. These results can be used to create visualizations summarizing the fluorescence in each brain region (Fig. 4c). The reported regions volumes are in voxel units, so the physical volumes will depend on the voxel dimensions used during imaging.

## Acknowledgements

We thank the entire Chung laboratory for support and discussions. K.C. was supported by the Burroughs Wellcome Fund Career Awards at the Scientific Interface, Searle Scholars Program, Packard award in Science and Engineering, NARSAD Young Investigator Award, and the McKight Foundation Technology Award. This work was supported by the JPB Foundation (PIIF and PNDRF), the NCSOFT Cultural Foundation, the Institute for Basic Science IBS-R026-D1, and the NIH (1-DP2-ES027992, U01MH117072)

## Author Contributions

J.S. worked on pystripe and prepared the main figures. L.K. worked on TSV and nuggt. K.X, Y.G.P. and D.H.Y prepared the example brain sample, and N.E. acquired the example data. G.D., K.X. and N.E. tested the protocol and provided feedback. J.S. and L.K. wrote the paper together, and all authors reviewed the manuscript. K.C. supervised the project.

---

**Supplementary Video 1. Destriping of NPY and SST image stack using pystripe.** Side-by-side flythrough of composite image stacks of NPY and SST before and after destriping using pystripe. The destriped stack was also corrected using illumination correction with the example flat reference image.

**Supplementary Video 2. 3D render of the stitched volumetric composite image.** Animated 3D render of syto 16, NPY, and SST composite volumetric images after destriping and stitching. Stitched images were downsampled by a factor of 2 to make the animation.

